# TrackRefiner: A tool for refinement of bacillus cell tracking data

**DOI:** 10.1101/2025.02.13.637647

**Authors:** Atiyeh Ahmadi, Alireza Dostmohammadi, Rhyan McLean, Brian Ingalls

## Abstract

**Motivation:** Single-cell resolution time-lapse microscopy of bacterial populations is a powerful tool for assessing cellular behavior and interaction dynamics. Realizing the full potential of this approach requires accurate image analysis: segmentation of individual cell objects, tracking of persistent cells from frame to frame, and connecting of mother cells to daughters when division events occur. In particular, accurate tracking is needed to produce longitudinal datasets for analysis of interactions and features that develop through time. Tracking is challenging when populations are densely packed or when cells undergo significant motion between frames. The leading software packages struggle to provide accurate data in such cases.

**Result:** To address this problem, we present TrackRefiner, a tool for refinement of bacillus cell tracking data. This package was specifically designed to refine the tracking outputs of CellProfiler, a commonly used image processing tool. TrackRefiner is built with a modular and publicly accessible structure, making it adaptable for integration with other image processing software. To assess the package’s performance, we manually determined ground truth tracking results for eight datasets from four research groups, comprising a total of 159,349 object links. This curated dataset, the first of its kind, serves as a valuable benchmark for assessing the performance of bacillus cell tracking algorithms. For timelapses with frequent imaging, TrackRefiner consistently achieved, with one exception, over 98% detection accuracy and corrected 57-100% of tracking errors. Accuracy was reduced for images sampled at lower frequency.

**Availability and implementation:** For easy access, TrackRefiner has been published on PyPI and Anaconda. Source code and user manuals can be accessed via Github and OSF. The manually curated benchmark dataset is also posted at these sites.

## Introduction

Time-lapse microscopy can noninvasively capture dynamic bacterial behavior. It has been used to study, for instance, how bacteria adapt to new environments, how they interact with other cells, and the development of resistance to antibiotics [4, 22, 8, 15, 29, 24]. In synthetic biology, time-lapse microscopy has been used to validate the behavior of engineered bacteria, making it a valuable tool for design and innovation [20, 9].

Interpretation of time-lapse microscopy observations typically requires the use of image processing tools to extract insight and understanding from large quantities of imaging data. When time-lapse images are captured at single-cell resolution (fields of view typically 100–500 µm, resolving cells measuring 1- 5 µm) tracking is needed to follow individual cells and their lineages. By tracking individual bacterial lineages, researchers can, for instance, pinpoint how antibiotic resistance evolves and propagates across successive generations [5, 3, 18], monitor metabolic shifts and gene expression regulation within specific lineages [13], and identify the emergence of specialized subpopulations, such as persisters that can withstand stress or antimicrobial treatments [19, 14]. Understanding these lineage-specific dynamics is important for devising strategies to control bacterial pathogens, improve biotechnological processes, and deepen our knowledge of microbial ecology and evolution.

The two main tasks in image processing are (i) object (cell) segmentation, i.e. distinguishing individual objects from the background and their neighbors, and (ii) object tracking, including tracking of cell division events [16]. Despite continued advancements in microscopy and image processing techniques, significant challenges remain.

Image processing approaches for non-motile bacillus populations were assessed in [1], which provided a benchmarked performance comparison of packages based on data from cell populations (of up to nearly a thousand cells) growing in constrained and unconstrained environments. That analysis showed that while currently available tools can deliver suitable segmentation performance in certain conditions, particularly via the recommended Omnipose-CellProfiler pipeline [11, 25], tracking performance is less satisfactory.

Ongoing efforts to improve segmentation performance are focused on developing advanced algorithms, including deep learning and hybrid models, to enhance accuracy and reliability across diverse imaging conditions [2, 11]. Accuracy of cell tracking depends on the accuracy of cell segmentation. However, even with perfect segmentation, tracking performance is inherently constrained by imaging frequency. More frequent imaging improves accuracy by reducing movement between frames, but practical limitations such as photobleaching, phototoxicity, and hardware constraints often prevent high-frequency imaging. Beyond these challenges, tracking of expanding cellular populations is inherently more complex than conventional computer vision tasks like vehicle tracking [21] due to the appearance of new (daughter cell) objects that must be correctly associated with their predecessors.

We thus developed TrackRefiner, a post-processing tool that takes segmentation and tracking output as a starting point and improves tracking accuracy by pruning incorrect links and adding missing links. TrackRefiner is a hybrid platform that combines traditional tracking methods with machine learning classifiers. It requires minimal user input, compensating for a lack of user-tuning by employing a data-driven approach. TrackRefiner was developed using test datsets of non-motile *Escherichia coli* and *Pseudomonas putida* populations cultured under agar pads or within microfluidic systems; it is generally applicable to bacillus bacteria.

TrackRefiner takes as input the processed output files from Omnipose-CellProfiler [25, 11]. (Adjustment to other package outputs would be straightforward, but is not provided in the current version; detailed adjustment instructions are provided on the TrackRefiner Developer Wiki). Specifically, it takes Omnipose-generated files (from the segmentation task), which include object masks, and CellProfiler’s tracking data, which includes a characterization of object (cell) features and spatial neighbor relations. The only user input required by TrackRefiner is the imaging interval time and the doubling time for the population. For scenarios in which cells leave the frame (e.g., exiting through the mouth of a microfluidic trap), the user can define a region of interest in which such events (termination of lineage tree branches) are not treated as tracking errors.

TrackRefiner’s output includes a pickle file for each time step storing tracking data and object feature lists, a CSV file with all tracking information, and a text log file providing details (time step, object, links) on error identification and correction and information about errors that were identified but not corrected (Additionally, we provide a separate GUI that allows users to locate and manually correct these errors.

To benchmark TrackRefiner’s performance, we manually generated ground truth tracking records for eight time-lapse experiments (contributed by four separate research groups [28, 12, 17]) and three representative snapshots (Table 1) capturing *E*.*coli* and *P*.*putida* populations cultured under agar pads or within microfluidic systems. The snapshots consist of pairs of timepoints for which the ratio of sampling frequency to growth rate is lower. The first and last frame of each dataset are shown in Supplementary Figure 1. This extensive ground truth dataset includes a total of 159,349 manually curated object links. We expect this dataset will be useful to the community as a benchmark for the ongoing assessment of tracking packages.

**Table 1.** Ground truth datasets

| Name | # Links | Average Doubling Time (min) | Imaging Interval (min) | Cell Type | # Frames |
| --- | --- | --- | --- | --- | --- |
| Unconstrained1 | 43156 | 30 | 2 | *P.putida* | 71 |
| Unconstrained2 | 7507 | 30 | 2 | *P.putida* | 23 |
| Unconstrained3 | 19903 | 30 | 2 | *P.putida* | 36 |
| Unconstrained4 | 1752 | 35 | 3 | *E.coli* | 71 |
| Constrained1 | 8615 | 20 | 10 | *E.coli* | 131 |
| Constrained2 | 5218 | 30 | 5 | *E.coli* | 40 |
| Constrained3 | 33751 | 45 | 1.75 | *E.coli* | 35 |
| Constrained4 | 33313 | 45 | 1.75 | *E.coli* | 35 |
| ConstrainedSnapshot1 | 304 | 20 | 5 | *E.coli* | 2 |
| ConstrainedSnapshot2 | 736 | 21 | 5 | *E.coli* | 2 |
| ConstrainedSnapshot3 | 732 | 22 | 5 | *E.coli* | 2 |

## Methods

The general pipeline of TrackRefiner is shown in Figure 1. TrackRefiner begins by making an assessment of segmentation based on object size. As described in detail below, cell object features are then collected from Omnipose- CellProfiler, followed by calculation of additional features. The algorithm then adjusts the tracking results by comparison with established biological patterns and dataset-specific observations, using both traditional assessments and machine learning classifiers. Finally, output files are produced.

**Fig 1.**
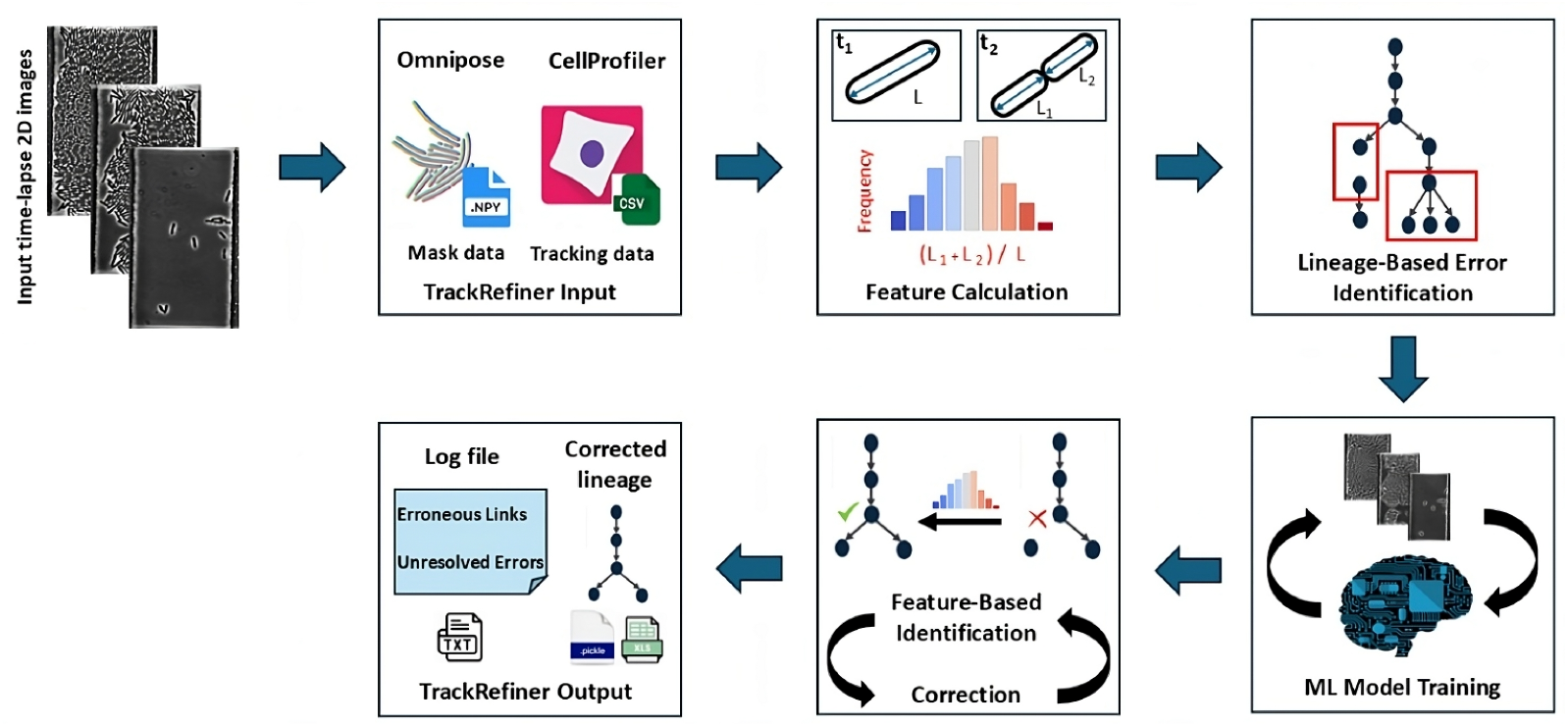
The TrackRefiner pipeline. The input consists of mask data (from Omnipose) and tracking, feature, and neighbor data (from CellProfiler). Additional features are then determined, after which errors revealed by lineage inconsistencies are identified. A set of machine learning models are then trained to distinguish correct and erroneous links. These trained models are then used for further error identification and for error correction. Output files provide corrected link information.

### Feature List

Tracking relies on an assumption that frame-to-frame changes are small, so a source object and a linked target object in the subsequent frame share many similarities. Here we describe the set of features that TrackRefiner uses for assessing and correcting track and division links. Some of these features are drawn directly from the CellProfiler output.

#### Cell Object Geometry

CellProfiler provides foundational cell object features such as length, orientation and centroid position. TrackRefiner calculates additional features capturing i) frame-to-frame elongation, ii) variation in length and elongation rate over an individual cell’s lifetime, iii) the difference in orientation between a mother cell and the average orientation of its daughters, and iv) the difference between a mother’s length and the lengths (sum and maximum) of its daughters.

#### Position

Omnipose provides a mask describing each cell object’s position [11]. TrackRefiner assesses consistency in position across frames via: i) the Intersection over Union (IoU) [16] (comparing mother and union of daughters in the case of division links) and ii) a proximity index determined by comparing the position of cell centroids and poles.

#### Neighbor Set

CellProfiler determines neighbors by detecting physical contact between expanded cell boundaries. TrackRefiner assesses changes in the neighbor count by considering neighbors that persist across consecutive frames. For division links, the neighbor set of the mother cell is compared to the union of the daughter cells’ neighbor sets.

#### Direction of Motion

TrackRefiner defines direction of motion by the vector between the centroids of the source and target cells (or from mother to each daughter). Additionally, the algorithm determines the alignment between a cell’s direction of motion and the average direction of motion of the neighbors of both the source and target cell.

### Algorithm Workflow

TrackRefiner uses a blend of feature analysis, machine learning classifiers, and systematic correction steps to address tracking challenges, as classified in Figure 1 (see also [27]).

In its first step, TrackRefiner identifies errors in segmentation based on a size threshold (i.e. non-cell objects that correspond to debris or optical “noise”). These objects and associated links are removed from the dataset. (This is the only way in which segmentation is assessed or corrected.)

TrackRefiner then collects and determines features across the remaining dataset. For some features, outlier ranges are determined by 95% confidence intervals via the normal approximation method [7].

### Error flagging

The algorithm then flags errors that are identifiable through anomalies in the lineage trees, specifically: (i) cells that branch into more than two daughter nodes (over-assigned division links), (ii) cell lineage origins after the first frame (untimely root) and (iii) cell lineage terminations before the last time frame (untimely leaf). Depending on context, these root and leaf events could be accurate descriptions of migration or death events. To address migration (cells leaving or entering the image frame), the user can specify regions in which such errors are not assigned (e.g. the mouth of a microfluidic trap). Cell death is not explicitly treated (but is the subject of planned future development.)

In addition to these lineage-based flags, TrackRefiner uses features to identify cells mistakenly treated as having divided when they did not (redundant division links) by comparing the length of the putative mother cell to the lengths of the putative daughters.

### Classifier training

After flagging readily identifiable errors in the previous step, the algorithm leverages CellProfiler link data that has not been flagged as erroneous (henceforth referred to as ‘verified’ links) to train a set of three classifiers. This ‘verified’ training set will generally contain errors, but is largely correct (more accurate than the pre-flagged errors reported in Table 2 below), and so is a solid foundation for a data-driven approach.

**Table 2.** TrackRefiner performance across the datasets in Table 1. Columns represent: the total number of links assessed in the ground truth (G.T.); the link error rate in Omnipose-CellProfiler output (false positives and false negatives); the number of segmentation errors (noise objects; N.O.) removed by TrackRefiner; the rate at which TrackRefiner identified tracking errors (the number of errors identified relative to the total error count); the TrackRefiner correction rate (net improvment in error count relative to the total error count); and the execution time.

| Dataset | Links in G.T. | Error(%) | N.O. Detected | Error Detection(%) | Error Correction(%) | Execution time (min) |
| --- | --- | --- | --- | --- | --- | --- |
| Unconstrained1 | 43,156 | 10.9 | 32 | 100.0 | 69.4 | 22.000 ± 0.010 |
| Unconstrained2 | 7,507 | 11.4 | 422 | 99.657 | 98.4 | 7.000 ± 0.030 |
| Unconstrained3 | 19,903 | 10.6 | 711 | 99.907 | 97.3 | 12.000 ± 0.020 |
| Unconstrained4 | 1,752 | 1.1 | 0 | 100.0 | 100.0 | 0.300 ± 0.006 |
| Constrained1 | 8,615 | 4.6 | 0 | 80.916 | 57.2 | 1.000 ± 0.090 |
| Constrained2 | 5,218 | 2.6 | 335 | 99.457 | 98.3 | 2.000 ± 0.070 |
| Constrained3 | 33,751 | 4.2 | 988 | 98.998 | 97.4 | 4.000 ± 0.080 |
| Constrained4 | 33,313 | 3.5 | 735 | 98.612 | 70.1 | 5.000 ± 0.060 |
| ConstrainedSnapshot1 | 265 | 23.0 | 0 | 49.18 | 39.34 | — |
| ConstrainedSnapshot2 | 510 | 32.3 | 4 | 44.84 | 44.24 | — |
| ConstrainedSnapshot3 | 573 | 31.0 | 2 | 42.37 | 32.76 | — |

Three separate classifiers are constructed, as described below. Details on classifier training and testing are provided in GitHub and OSF.

**Classifiers:** The algorithm’s **primary classifier** assesses a putative link (i.e. a source-target pair) and produces two likelihoods: (i) that the pair is connected by a track link, and (ii) that the pair is connected by a division link. Because these likelihoods are produced from a single classifier, they can be usefully compared. The positive sets are verified links. The negative sets are verified links of opposite type.

In addition to the primary classifier, the algorithm trains two **specialist classifiers**: one for assessing putative track links and another for assessing putative division links. These are, in a sense, redundant, because the primary classifier produces corresponding likelihoods. However, these specialist classifiers each focus exclusively on one link type, allowing them to refine predictions beyond the primary classifier’s general approach. Consequently, they individually provide more confident estimates of the likelihood of link accuracy. For the specialist division link classifier, the positive class is verified division links; the negative class is verified track links. For the specialist track link classifier, the positive class is verified track links; the negative class is verified division links and additional fabricated track links that connect sources of verified track links to neighboring cells of the target (as additional examples of links that should be recognized as inaccurate).

### Correction phase

After training the three classifiers, TrackRefiner uses them to resolve tracking errors flagged in the initial identification step and to address feature-based errors as described below. For over-assigned division links, the specialist division link classifier is applied to retain the two most likely division links. Redundant division links are addressed in a similar manner using the specialist track link classifier. Swapped track links and missing division links, which are not easily identifiable through lineage tree anomalies, are identified based on feature-based discrepancies, such as deviations in motion alignment, neighbor ratios, and length dynamics. These potential errors are evaluated and corrected using the primary comparison classifier. For untimely root and leaf errors, TrackRefiner applies the primary comparison classifier to assess nearby candidates and re-establish probable links using the Linear Sum Assignment Method (LSAM) [10].

**Fig 2.**
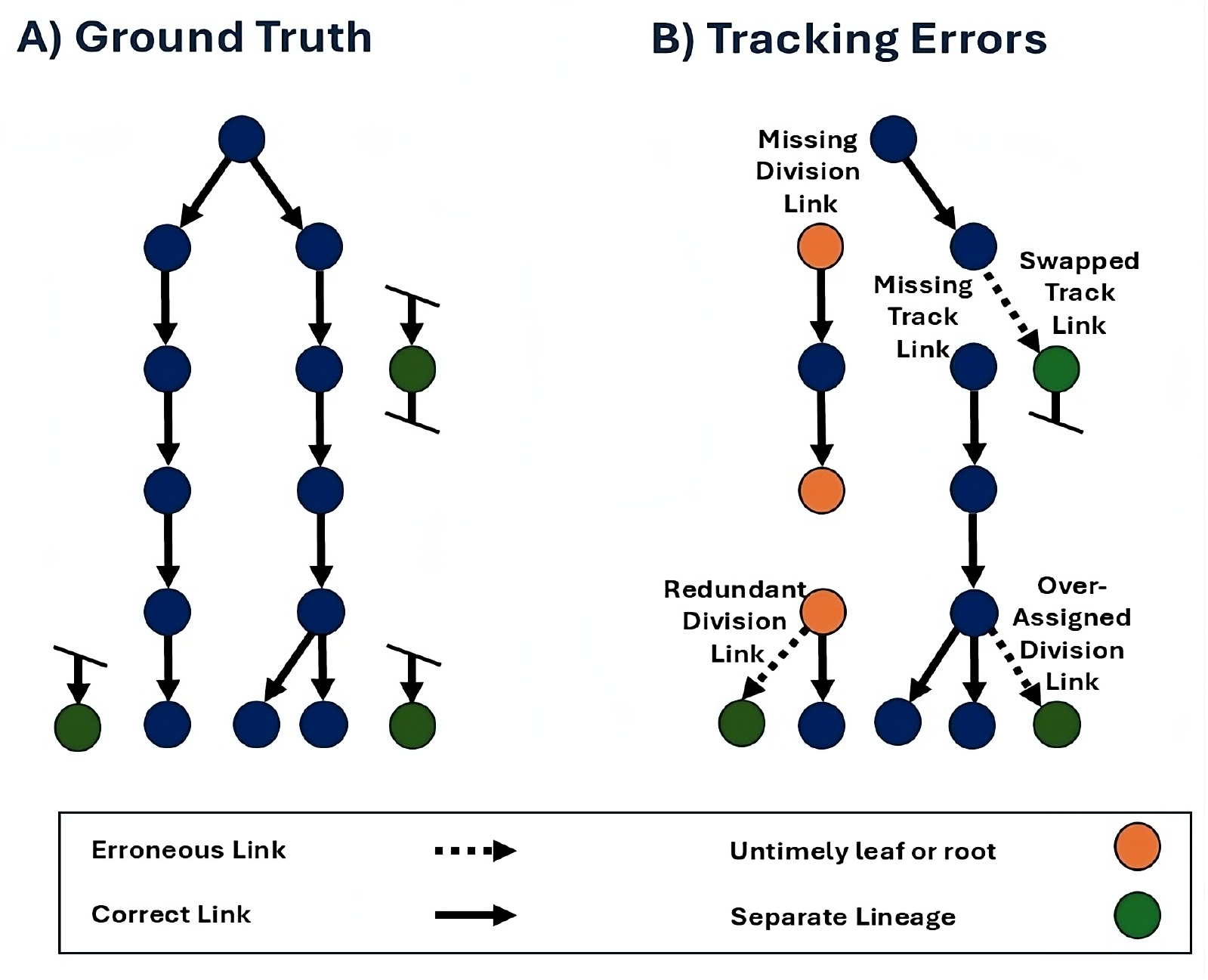
Classification of tracking errors. (A) Ground truth showing true track and division links. (B) Tracking errors: missing track and division links, over-assigned division links, redundant division links, swapped track links, and untimely root or leaf assignments (i.e. lineage start/end points that are not in the initial/final frame).

Any unresolved untimely root or leaf cases are revisited in a final restoring step in which the primary classifier re-evaluates link probabilities and establishs connections where it deems appropriate. The correction process successfully resolves many errors, but it can also introduce erroneous links. Algorithm details are provided in GitHub and OSF.

### Ground truth preparation

To evaluate the accuracy of TrackRefiner, we manually generated ground truth benchmarks for the datasets described in Table 1 (see also Supplementary Figure 1). Each link reported by CellProfiler was manually reviewed and labeled as correct or erroneous; simultaneously, missing links were identified and labeled. This process produced a comprehensive assessment of Omnipose-CellProfiler errors and a ground truth consisting of 159,349 labeled links across a total of 442 frames. The same comparison process was used to assess errors in TrackRefiner output. We also applied this procedure to the snapshot datasets (Table 1; Supplementary Figure 1), assessing 1771 links over 6 frames.

A single team member manually identified all tracking errors. This assessment was independently reviewed by a team of three different individuals. To facilitate this process, we developed two specialized GUIs (for CellProfiler output and for TrackRefiner output).

## Results

Table 2 summarizes TrackRefiner’s performance, showing dataset size, the rate of inaccurate or missing link reports, the number of segmentation errors (i.e. noise objects) removed in TrackRefiner’s first step, and TrackRefiner’s primary performance: how effectively TrackRefiner identifies tracking errors (reported as the number of errors identified relative to Omnipose-CellProfiler’s total error count) and the improvement in accuracy (reported as the net improvement in error count relative to Omnipose-CellProfiler’s total error count). This correction rate is bounded above by the error identification rate. Execution time is also reported (not included for the snapshots). All analyses were performed on a system equipped with an Intel(R) Core™ i9-13900 CPU, 128 GB of RAM, and a 1 TB NVMe SSD.

For the full datasets (Unconstrained1-4, Constrained1- 4), TrackRefiner consistently achieved high detection rates, above 98%, with one exception, and corrected a substantial proportion of errors, ranging from 57% to 100% of the total error count.

The snapshot datasets exhibit a higher rate of tracking errors: in these scenarios the ratio of imaging frequency to division rate is low (Table 1), so significant motion can occur between frames. Moreover, cells in these frames lack consistent alignment, resulting in erratic trajectories and frequent changes in orientation. For these cases, the TrackRefiner error detection rate drops (ranging from 42% to 49%), with reduced error correction rates (32% to 44% of the total error count).

To assess TrackRefiner’s capacity to deliver improvements in longitudinal analysis, we determined TrackRefiner’s performance in correcting cell cycle duration results (Figure 3) across the eight complete datasets. TrackRefiner reduced errors compared to Omnipose-CellProfiler results (details available in GitHub).

**Fig 3.**
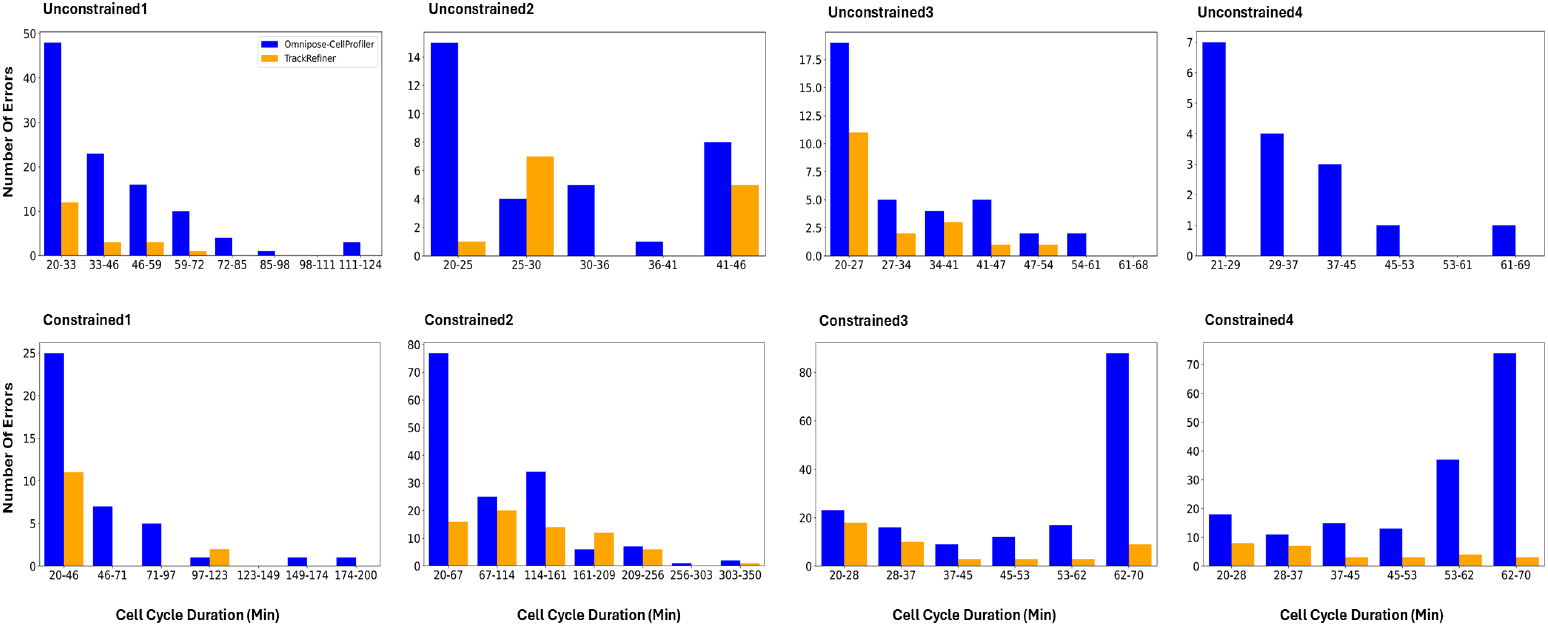
TrackRefiner reduces error in assessment of cell cycle duration. For each dataset, histogram bins were selected to describe the distribution of cell cycle durations in the ground truth (shown in GitHub). The distribution of cell cycle durations reported by Omnipose-CellProfiler and TrackRefiner were assigned to the same bins. The difference (absolute error) in bin heights is shown.

### Considerations for Tracking Accuracy

Tracking performance is affected by a variety of biological and technical factors, and depends on both the spatial arrangement of cells and the temporal dynamics of their movement (which is directly dependent on imaging frequency). We used the ground truth dataset to investigate these relationships, as follows.

Figure 4A shows the relationship between average absolute angular motion and the percentage of tracking errors made by Omnipose-CellProfiler; Figure 4B shows the same data for average absolute translational motion. In all cases for which there is sufficient data to confirm a trend, increased motion leads to increased error rate. A similar analysis of the effect of density yielded results that are more ambiguous (Supplementary Figure 2). Although dense packing increases consistency in motion, it also increases the segmentation error rate; our analysis is confounded by this combination.

**Fig 4.**
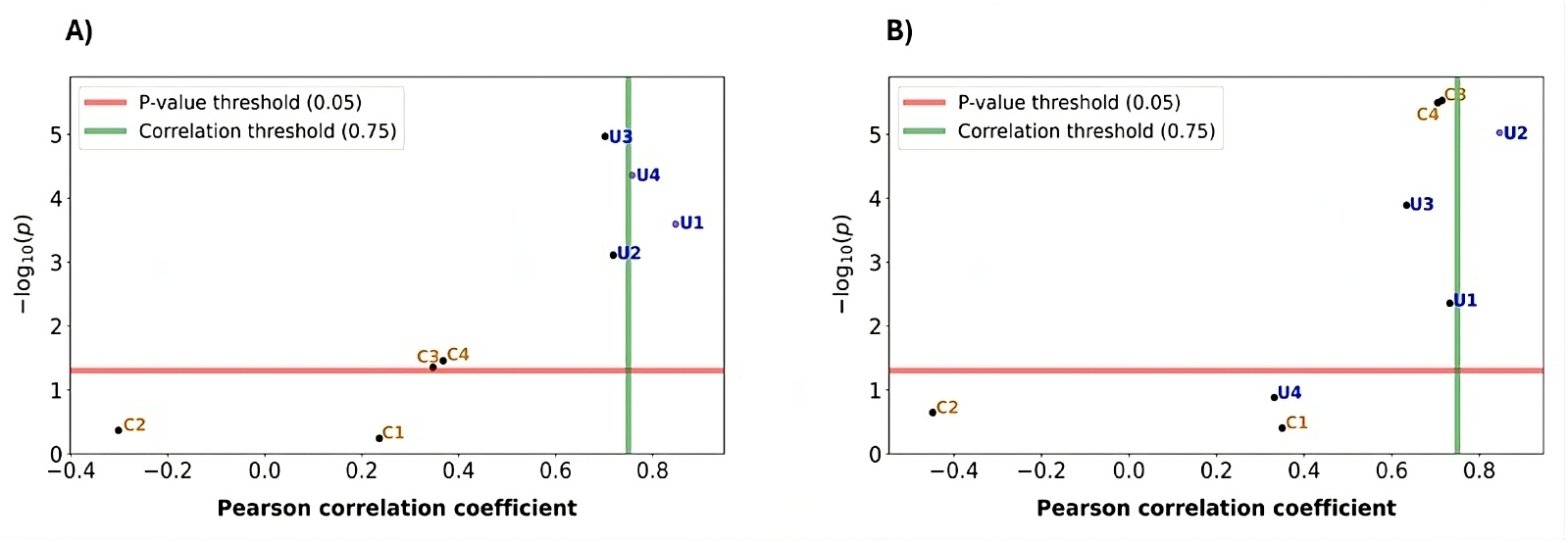
Correlation of angular (A) and translational (B) motion with Omnipose-CellProfiler tracking error rate for each dataset. For each of the eight full time-lapses (U: unconstrained, C: constrained), the horizontal axis represents Pearson’s correlation coefficient (*r*) between the average motion (A: angular; B: translational of centroids) and the percentage of tracking errors. The vertical axis shows -log_10_(p), where *p* is the p-value calculated using a t-distribution (details in GitHub). In both panels, the horizontal red line marks *p* = 0.05 (-log_10_(p) ≈ 1.3), which serves as the threshold for statistical significance. The vertical green line at *r* = 0.75 indicates a threshold for a strong positive correlation.

## Discussion

TrackRefiner is a specialized software package designed to detect and refine tracking errors in the output of existing image processing algorithms. In its current implementation, it utilizes segmentation results from Omnipose and tracking data from CellProfiler. TrackRefiner combines traditional assessments with machine learning to analyze multiple features simultaneously, making it well-suited for large and complex datasets.

TrackRefiner is designed with accessibility in mind, requiring no prior knowledge of machine learning or programming. It operates as a plug-and-play system, where users input basic parameters (imaging interval time and cell doubling time). Furthermore, TrackRefiner offers both a command-line and Graphical User Interface, catering to users from computational biologists to experimentalists.

We evaluated TrackRefiner’s performance against a range of datasets using ground truth tracking data. This benchmark dataset, the first of its kind, provides a valuable resource for assessing future developments in tracking software. TrackRefiner consistently achieved high detection rates, above 98%, with one exception, and corrected a substantial proportion of errors, ranging from 57% to 100% of the total error count. For more challenging snapshot datasets, detection rates dropped below 50%, while correction accounts for 32% to 44% of the total error counts. TrackRefiner produces a text log file that provides detailed information on the errors that it has identified but has not corrected; these can be located and manually fixed with a specialized GUI).

While TrackRefiner addresses many challenges in cell tracking, challenges clearly remain. TrackRefiner’s machine learning performance benefits from larger datasets with more links to ensure effective learning of patterns. Moreover, its accuracy depends on the quality of segmentation from Omnipose-CellProfiler; segmentation errors propagate into the tracking phase, limiting overall performance. Our analysis was limited to time-lapse images of non-motile bacilli. Extension to other cellular morphologies and physiologies will be needed to ensure broader utility.

To further improve tracking accuracy, future iterations of TrackRefiner could benefit from integrating Graph Neural Networks (GNNs), which are highly effective at analyzing dynamic, interconnected data like time-lapse cell interactions [6]. Enhancing compatibility with additional tools, such as SuperSegger [26] and DelTA [23], would also increase its utility. A key challenge lies in mitigating motion-induced errors. Predictive modeling techniques that leverage historical trajectories and dynamic adaptation could help anticipate cell movements. Beyond improving individual tracking events, TrackRefiner contributes to a more fundamental goal: enabling researchers to reconstruct accurate single-cell histories, trace lineage relationships, and better understand microbial dynamics over time. As imaging technologies continue to advance, precise tracking will remain critical for bridging the gap between raw microscopy data and meaningful biological insights.

## Conclusion

TrackRefiner offers an effective solution for improving cell tracking in time-lapse microscopy of bacillus populations. By combining traditional methods with machine learning, it refines tracking outputs from Omnipose-CellProfiler. To rigorously assess its performance, we established a manually curated ground truth dataset, reviewing 159,349 individual tracking links across multiple time-lapse experiments. Our evaluation showed that TrackRefiner corrects a large portion of tracking errors while remaining easy to use with its simple interface and minimal setup requirements. TrackRefiner is designed to be open-source and completely transparent, making it accessible for both programmers and non-coders alike. A detailed online tutorial is available in TrackRefiner Wiki.

## Author contributions statement

Atiyeh Ahmadi: Conceptualization, algorithm design, development, and testing, ground truth creation, data processing, Writing – review and editing. Alireza Dostmohammadi: Code optimization, algorithm development. Rhyan McLean: Ground truth creation. Brian Ingalls: Conceptualization, Funding acquisition, Supervision, Writing – review and editing

## Acknowledgment

This work was supported by a Discovery Grant (RGPIN- 2018-03826) from Canada’s Natural Sciences and Engineering Research Council (NSERC). We extend our heartfelt thanks to Clara M. Heiman, Jordan Vacheron, Sander Tans, Simon van Vliet, and Aaron Yip for generously sharing their data, which played a crucial role in helping us create a comprehensive ground truth from multiple groups around the world. We would like to express our gratitude to the first users of TrackRefiner, Aaron Yip and Amélie Chandoboius, for their patience. We would also like to thank Samantha Schwartz for her assistance in reviewing the ground truth preparation. This work was carried out on the Haldimand Tract, land granted to the Haudenosaunee (Six Nations of the Grand River) in 1784. Settler colonial theft of this land is ongoing.

