## Supplement for "TrackRefiner: A tool for refinement of bacillus cell tracking data"

### Appendix

#### Datasets

Figure 1 shows the first and last image from of each dataset.

#### Density Plot

Figure 2 shows the correlation of pixel density with Omnipose-CellProfiler tracking error rate for each time-lapse (with the calculation of pixel density explained in [2]. The results are largely inconclusive.

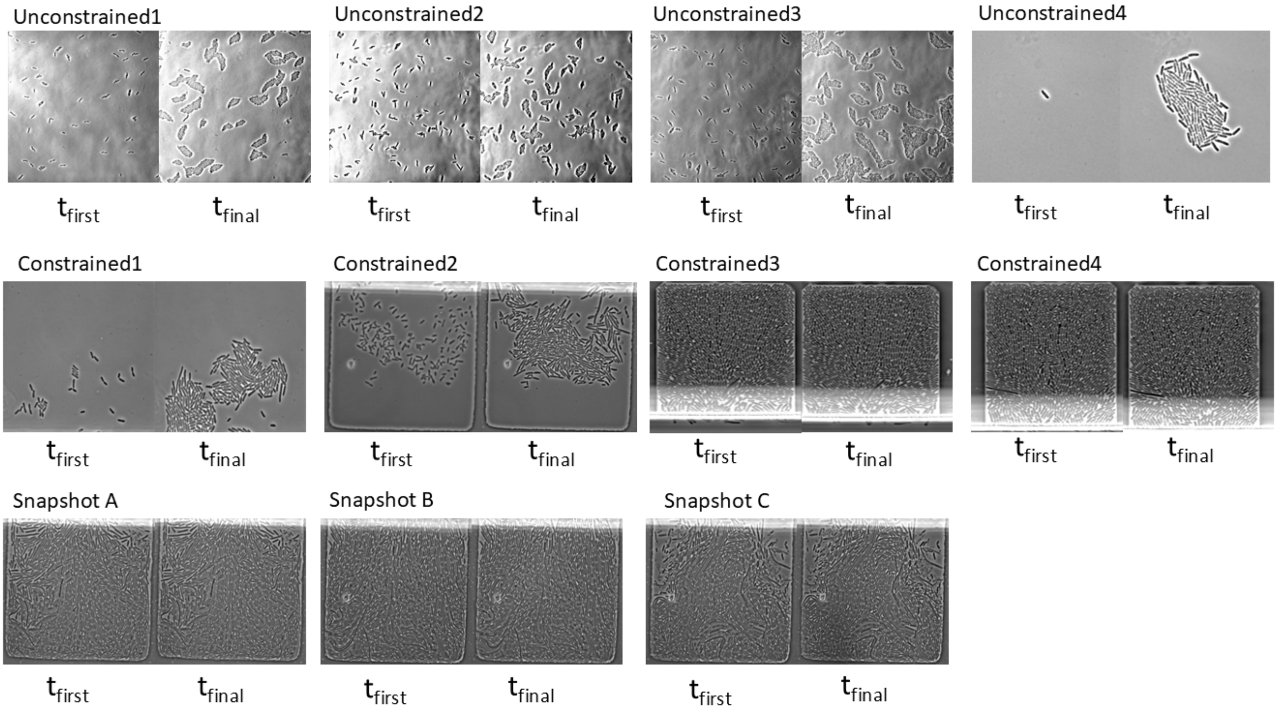


**Fig. 1. Dataset Samples.** The first and last time points of each dataset are shown. The images were preprocessed using the filters recommended in [1].

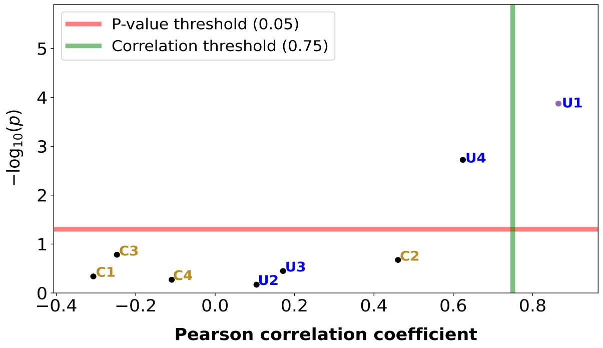


**Fig. 2. Correlation of pixel density with tracking error rate for each time-lapse.** For each of the eight full time-lapses (U: unconstrained, C: constrained), the horizontal axis represents Pearson's correlation coefficient ($r$) between pixel density and the percentage of tracking errors. The vertical axis shows -$\log_{10}$(p), where $p$ is the p-value calculated using a t-distribution (details in GitHub). In the panel, the horizontal red line marks $p = 0.05$ (-$\log_{10}$(p) $\approx$ 1.3), which serves as the threshold for statistical significance. The vertical green line at $r = 0.75$ indicates a threshold for a strong positive correlation.
